# Lanthanide-dependent alcohol dehydrogenases require an essential aspartate residue for metal coordination and function

**DOI:** 10.1101/2020.02.25.964783

**Authors:** Nathan M. Good, Matthias Fellner, Kemal Demirer, Jian Hu, Robert P. Hausinger, N. Cecilia Martinez-Gomez

**Affiliations:** Department of Microbiology & Molecular Genetics, Michigan State University, East Lansing, MI 48824; Department of Biochemistry, Michigan State University, East Lansing, MI 48824; Biochemistry, University of Otago, P.O. Box 56, Dunedin, Otago, 9054, New Zealand; Okemos High School, Okemos MI 48864; Department of Chemistry, Michigan State University, East Lansing, MI 48824

**Author notes:** Both authors contributed equally to the work presented (Nathan M. Good, Matthias Fellner). To whom correspondence should be addressed: N. Cecilia Martinez-Gomez: Department of Microbiology & Molecular Genetics, Michigan State University, East Lansing 48824.

**Keywords:** XoxF, ExaF, lanthanide, methanol dehydrogenase, ethanol dehydrogenase, alcohol dehydrogenase, metalloprotein, pyrroloquinoline quinone, cofactor coordination, crystallography

## Abstract

The presence of lanthanide elements (Ln^3+^) and pyrroloquinoline quinone (PQQ) containing cofactors in XoxF methanol dehydrogenases (MDHs) and ExaF ethanol dehydrogenases (EDHs) has expanded the list of biological elements and opened novel areas of metabolism and ecology. Other MDHs known as MxaFIs are related in sequence and structure to these proteins, yet they instead possess a Ca^2+^-PQQ cofactor. An important missing piece of the Ln^3+^ puzzle is defining what protein features distinguish enzymes using Ln^3+^-PQQ cofactors from those that do not. In this study, we investigated the functional importance of a proposed lanthanide-coordinating aspartate using XoxF1 MDH from the model methylotrophic bacterium *Methylorubrum extorquens* AM1. We report two crystal structures of XoxF1, one containing PQQ and the other free of this organic molecule, both with La^3+^ bound in the active site region and coordinated by Asp320. Using constructs to produce either recombinant XoxF1 or its D320A variant, we show Asp320 is needed for *in vivo* catalytic function, *in vitro* activity of purified enzyme, and coordination of La^3+^. XoxF1 and XoxF1 D320A, when produced in the absence of La^3+^, coordinate Ca^2+^, but exhibit little or no catalytic activity. In addition, we generated the parallel substitution to produce ExaF D319S, and showed the enzyme loses the capacity for efficient ethanol oxidation with La^3+^. These results provide empirical evidence of an essential Ln^3+^-coordinating aspartate for the function of XoxF MDHs and ExaF EDHs; thus, supporting the suggestion that sequences of these enzymes, and the genes that encode them, are markers for Ln^3+^ metabolism.

---

Lanthanide elements (Ln^3+^) have been shown to greatly impact methylotrophy—the ability of microorganisms to derive all carbon and energy needed for survival and growth from reduced compounds lacking carbon-carbon bonds, such as methane and methanol (1–5). Ln^3+^ associate with pyrroloquinoline quinone (PQQ) and function as cofactors of XoxF-type methanol dehydrogenases (MDHs) and ExaF-type ethanol dehydrogenases (EDHs) in methylotrophic bacteria (6–9). These enzymes are aptly referred to as Ln-dependent alcohol dehydrogenases (Ln-ADHs) and their discovery has added Ln^3+^ to the biological table of the elements (7, 8, 10–12). A number of XoxF-type MDHs have been studied from methylotrophic bacteria, with analyses including X-ray crystallography and enzyme kinetics (7, 9, 13–16). ExaF EDH from the model methylotroph *Methylorubrum* (formerly *Methylobacterium* (17)) *extorquens* AM1 is currently the only representative reported from a methylotroph, though genes encoding putative ExaF homologs have been identified in a diverse set of organisms (8, 18, 19). Phylogenetic analyses of XoxF-coding genes indicate they are wide-spread in the environment and can be grouped into at least 5 distinct clades with representatives in *Alpha-, Beta-*, and *Gammaproteobacteria*; *Verrucomicrobia*, and the NC10 phylum bacterium *Candidatus* “Methylomirabilis oxyfera” (20). Importantly, phylogenetic consideration of potential Ln^3+^-related genes led to two discoveries: 1 - bacteria previously reported to be non-methylotrophic, such as *Bradyrhizobium diazoefficiens* USDA110, can indeed grow methylotrophically using XoxF-type MDHs (15), and 2 - Ln-ADHs metabolize multi-carbon compounds in methylotrophs and non-methylotrophs, such as *Pseudomonas putida* KT2440 (8, 21). These relatively recent discoveries underscore the relevance of Ln^3+^ to microbial diversity and emphasize the importance of metal bioavailability on plant, soil, aquatic, and marine ecosystems—complex environments where Ln-utilizing bacteria are major constituents (22–25).

Lighter versions of Ln^3+^, ranging from lanthanum (La^3+^) to europium (Eu^3+^) (atomic numbers 57-63), excluding promethium (Pm^3+^), have been shown to function with PQQ as essential cofactors in XoxF-type and ExaF-type ADHs. Ln-ADHs can be distinguished from MxaFI-type MDHs and ExaA-type EDHs that bind PQQ and coordinate Ca^2+^ (26, 27). In the heterotetrameric MxaFI MDHs, Ca^2+^ serves as a Lewis acid that polarizes the C5 carbonyl of PQQ, facilitating hydride transfer from the alcohol substrate (Fig. 1*A*) (28). Because the active site of ExaA-type EDHs is very similar to that of MxaFI, the reaction mechanism is likely analogous (29). Initial reports of Ln-ADHs showing higher catalytic efficiencies compared to Ca-ADHs generated excitement that Ln^3+^ coordination augmented ADH efficiency as a general phenomenon (7, 8, 20). However, characterization of additional Ln-ADHs included some exhibiting catalytic efficiencies similar to Ca-ADHs (6–9, 13, 14, 20, 21). It is possible that unique physiologies of certain bacteria, such as thermoacidophiles like *M. fumariolicum* SolV, require extremely efficient XoxF MDHs for survival (7), but this is likely the exception and not the rule for Ln-ADHs. Functional redundancy may also allow for the adaptation of secondary enzymes for alternative substrates with distinct roles in metabolism, such as formaldehyde oxidation by ExaF (8, 13).

**FIGURE 1.**
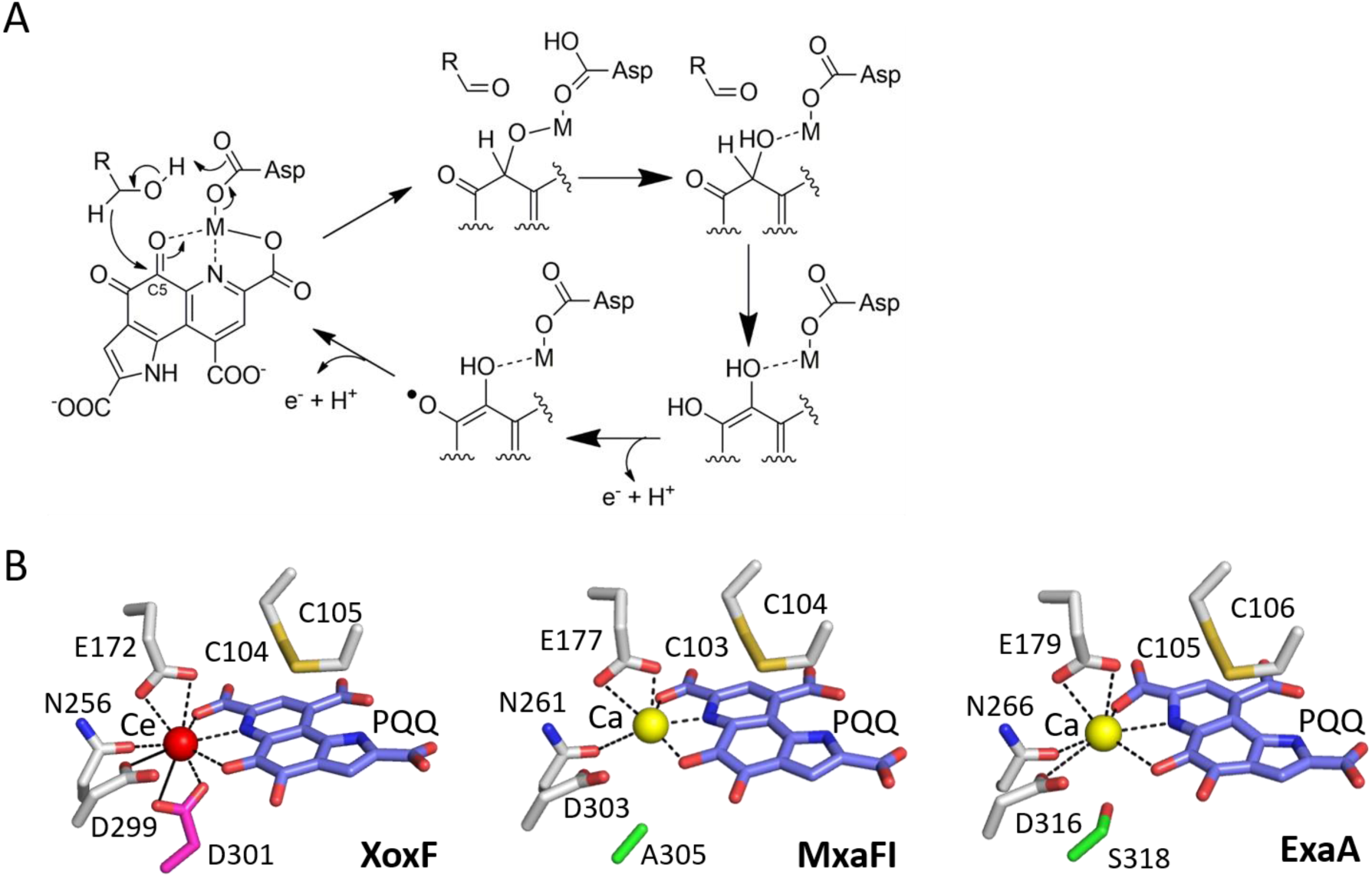
Catalysis of alcohol oxidation by PQQ ADHs. *A*, Catalytic mechanism of alcohol oxidation. The reactive C5 carbonyl of PQQ and Asp required for catalysis are indicated. *B*, Active site structures of PQQ ADHs: XoxF from *M. fumariolicum* SolV (PDB ID: 4MAE) (7), MxaFI from *M. extorquens* AM1 (PDB ID: 1W6S) (80) and ExaF from *P. aeruginosa* (PDB 1FLG) (81). The Asp residue depicted in the mechanism (*A*) corresponds to D299, D303, and D316 in the three structures (*B*). PQQ, slate; cerium, red sphere; calcium, yellow sphere; adjacent Cys residues that form a characteristic disulfide, orange; conserved Asp in Ln ADH, hot pink (XoxF); Ala (MxaFI) or Ser (ExaA) in the corresponding position in Ca ADH, green.

Although Ln-ADHs and Ca-ADHs share many similarities, their clear differences in metal preference raise the question of what structural features, if any, determine and are potentially diagnostic for metal usage. XoxF-type MDHs are α_2_ homodimeric enzymes and distinct from the α_2_β_2_ heterodimeric structure of MxaFI-type MDHs (30, 31). One important exception to this generalization is XoxF-type MDH from *Candidatus* “Methylomirabilis oxyfera”, which was purified as a α_2_β_2_ heterodimer that included MxaI, the small-subunit of MxaFI-type MDHs (16). ExaF-type and ExaA-type EDHs are both α_2_ homodimeric enzymes (8, 32). Quaternary structure alone, therefore, cannot be used to differentiate Ln-ADHs and Ca-ADHs. Multiple sequence alignment of type I ADHs, which includes both MDHs and EDHs, shows high conservation of catalytically and structurally important amino acids, including Asn, Glu, and Asp residues in the active site (20). Remarkably, one residue is differentially conserved in Ln-ADHs compared to Ca-ADHs (Fig 1*B*). Nearly all XoxF and ExaF sequences have an additional Asp residue positioned two amino acids downstream from a conserved Asp required for catalytic function (20). This position is occupied by Ala in MxaFI-type MDHs and Ser or Thr in ExaA-type EDHs. The crystal structure of XoxF MDH from *M. fumariolicum* SolV revealed this additional Asp to coordinate the Ce^3+^, and this residue was proposed to be diagnostic for Ln-ADHs (7, 20). The few XoxF-type MDH crystal structures generally support the role of this residue as being important for Ln^3+^ coordination and function of the enzyme (7–9, 33–35).

*M. extorquens* AM1 has been a model of study for one-carbon metabolism for decades. This methylotrophic bacterium produces XoxF1 MDH and ExaF EDH that both contain Ln-PQQ cofactors (5, 13, 36–38) and possess the additional Asp residue proposed to be important for Ln^3+^ coordination: Asp320 (XoxF1) and Asp319 (ExaF), respectively. XoxF1 and ExaF have been kinetically characterized with La^3+^, and XoxF1 with neodymium as well (8, 13). *M. extorquens* AM1 also produces the Ca^2+^-dependent MxaFI MDH. Expression of the *mxa* operon, encoding MxaFI and accessory/Ca^2+^-insertion proteins, is differentially regulated from the *xox1* gene cluster by the “Ln switch” that is sensitive to the presence of nanomolar Ln^3+^ (36, 39, 40). When Ln^3+^ are absent from the growth medium (or at sub-nanomolar concentrations), *xox1* expression is down-regulated and *mxa* expression is up-regulated. If Ln^3+^ are present at nanomolar or greater concentrations, *mxa* expression is down-regulated and *xox1* is highly expressed (13). The presence of the Ln switch in *M. extorquens* AM1 and the capacity to produce Ca-MDH and Ln-ADH make it an excellent model for the study of Ln biology.

Of the few reported Ln-ADHs, all have the hypothetical “Ln-coordinating Asp” including XoxF1 and ExaF from *M. extorquens* AM1. Although theoretical studies support the importance of this residue for the function of Ln-ADH, no empirical study, until now, has shown this residue is required for Ln^3+^ coordination and catalytic function of these enzymes. We report a 3.1 Å crystal structure of XoxF1 from *M. extorquens* AM1 as a representative Type V XoxF MDH structure. The protein crystallized as a homodimer with one La^3+^ and one PQQ per subunit. We report an additional 2.8 Å structure with only La^3+^ coordinated; this is the first structure of an MDH without PQQ bound to our knowledge. In both structures, Asp320 contributes to La^3+^ coordination. Using site-directed mutagenesis, we constructed an Ala320 substitution variant of XoxF1 (D320A) from *M. extorquens* AM1 and show the mutant cells were incapable of growth with methanol and La^3+^. MDH activity was only detectable for XoxF1, and not for XoxF1 D320A, when purified from cultures grown with La^3+^. When enriched from cultures lacking La^3+^, XoxF1 and XoxF1 D320A exhibited little to no activity with methanol. Further, we show that production of catalytically inactive XoxF1 from plasmid constructs was sufficient to allow for MxaFI-dependent growth on methanol in a Δ*xoxF1* Δ*xoxF2* mutant. Finally, we report that an ExaF D319S variant is inactive with ethanol *in vivo*, providing evidence that the Ln-coordinating Asp is also necessary for catalytic function of ExaF-type EDHs. Overall, this study provides empirical evidence in support of the Ln-coordinating Asp being necessary for Ln-ADH catalytic function and supports its potential use as a marker to identify new Ln-ADHs.

## RESULTS

### C*rystal structures of XoxF1 with La*^*3+*^

The number of Ln-ADH crystal structures available is limited and more representatives are needed to better understand structural similarities and differences among Ln- and Ca-ADHs. Currently, only three structures are available for study: XoxF from *M. fumariolicum* SolV with Ce^3+^ or Eu^3+^ (PBD ID: 4MAE (7) or 6FKW (41)), and XoxF from *Methylomicrobium. buryatense* 5GB1C (PBD ID 6DAM; (9)). The enzyme from *M. fumariolicum* SolV falls within the type II clade of XoxF MDH, and that from *M. buryatense* is a type V enzyme. Both of these organisms are methanotrophs (capable of oxidizing methane to methanol), whereas *M. extorquens* AM1 cannot oxidize methane. *M. extorquens* AM1 has a type V enzyme, XoxF1, which was the first Ln-ADH described in the scientific literature (6). Using immobilized metal affinity chromatography (IMAC), we purified recombinant XoxF1 fused to a hexa-histidine tag from cultures grown in minimal methanol medium with 20 µM LaCl_3_. After tag cleavage and concentration to 2.5 mg/mL, the protein was crystallized (see methods). Two XoxF1 structures were resolved: one in complex with La^3+^-PQQ, and a second with only La^3+^ bound (Fig. 2). The overall structures, both showing two protein chains in each asymmetric unit, are nearly identical with a Cα alignment resulting in a root-mean-square deviation (RMSD) of ∼0.3 Å comparing chains across and within the two structures (Table S1). All chains are fully built from residue Asn22 to the penultimate C-terminal residue, Asn600. The missing N-terminal residues were previously identified as a likely signal peptide for translocation from the cytoplasm to the periplasm (20). The overall fold matches other MDHs with the typical eight-bladed β-sheet propeller surrounding the active site (Fig. 2*A*) (9, 20, 30, 31, 42–46). Comparison to the most closely related methanol dehydrogenases with available structures shows minor deviations in surface exposed loops: *M. buryatense 5G* XoxF (PDB ID 6DAM (9)) had ∼0.5 Å Cα RMSD with 67% sequence identity and *M. fumariolicum SolV* XoxF (PDB ID 4MAE (7) or 6FKW (41)) had ∼0.7 Å Cα RMSD with 55% sequence identity (Figure S1*A*).

**FIGURE 2.**
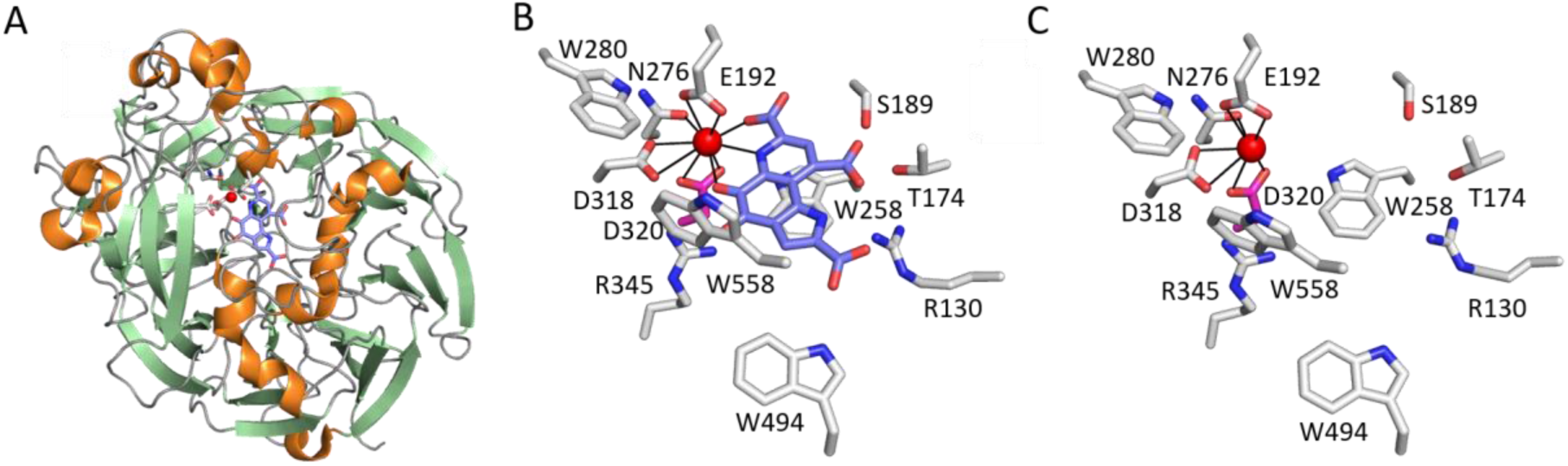
*M. extorquens* AM1 XoxF1 crystal structures. *A*, Overall fold (La^3+^-PQQ model shown, PDB ID 6OC6) with β-sheets in pale green, α-helices in orange, and coils in grey. The active site is illustrated with La^3+^ in red; chelating residues in white; PQQ in slate; oxygen atoms in red; nitrogen atoms in blue. *B*, XoxF1-La^3+^-PQQ active site with metal-coordinating residues labelled; Ln-coordinating Asp in hotpink. *C*, XoxF1-La^3+^ active site region. La^3+^ and residues shown are the same as depicted in *B*.

The active site regions of the XoxF1-La^3+^-PQQ (Figure 2*B*) and XoxF1-La^3+^ (Figure 2*C*) show high similarity in structures. La^3+^ is coordinated the same way in both proteins using Glu192 (bidentate), Asn276 (monodentate via oxygen), Asp318 (monodentate), and Asp320 (bidentate). PQQ introduces three additional coordination atoms (two oxygen and one nitrogen) for the first structure. Residues surrounding PQQ show similar side chain rotamers in both XoxF1-La^3+^-PQQ and XoxF1-La^3+^ structures. In the La^3+^ only bound structure, chain A is 100% occupied by the metal and one of the PQQ coordination sites is occupied by a small molecule that we interpreted conservatively as a water. Chain B appears to be more disordered in the active site region and the La^3+^ atoms refined to 61% occupancy, indicating 39% of the structure is the metal-free state. This situation led to greater mobility of Trp280, Asp318, and Arg345 sidechains, likely indicating alternative conformations. In addition, Trp258 possibly shows a second conformation pointing towards La^3+^ and partially occupying its space in the metal free portion of the protein; however, at an overall resolution of 2.8 Å the minor alternate state of the protein could not be modelled with confidence. Given the apparent flexibility of these four residues, they may play a role in metal binding and metal release even though they do not directly coordinate the La^3+^. Alternatively, these residues could passively fill the cavity when La^3+^ is not present. Notably, the conformation of Asp320 did not change with decreased metal occupancy indicating likely inflexibility at this position. We speculate that PQQ also was present in the XoxF1-La^3+^ sample during its purification, but that the crystal conditions (with 10% propanol) resulted in release of the organic portion of the cofactor in both chains and partial loss of La^3+^ in chain B. When we regrew these crystals, substituting propanol with 10% methanol, we again obtained the XoxF1-La^3+^ structure lacking PQQ (data not shown). The implications of these observations are that XoxF1 coordinates La^3+^ even though PQQ is no longer part of the cofactor complex.

To compare with existing structures, we compiled a list of related MDH structures by cross-referencing entries of Pfam family PQQ_2 (PF13360) (47) having 25% sequence identities to *M. extorquens* XoxF1 at rcsb.org (48), as well as 3D structure hits better than 2.1 Å RMSD in DALI (49). Nineteen structures were examined after excluding 6 structures that shared the overall fold, but not the general active site environment, and we found that all proteins had both the metal and PQQ bound in the active site. The XoxF1-La^3+^ structure reported here is currently the only PQQ-free structure of this family, and notably the “organic cofactor-less” enzyme maintained the homodimer quaternary structure. From eleven unique proteins, the XoxF1 La^3+^-PQQ active site environment is very similar to the two most closely related Ln-ADH (6DAM La^3+^-PQQ, 4MAE Ce^3+^-PQQ, 6FKW Eu^3+^-PQQ). In all cases the same protein side chain and PQQ metal chelation is observed, suggesting that light Ln^3+^ share the same coordination in this state (Figure S1*B*) as predicted by DFT calculations (33, 50). The nine remaining proteins have a Ca^2+^ atom (with one Mg^2+^ exception) bound in their structures. The main difference to XoxF1 from *M. extorquens* AM1 is seen in the position corresponding to Asp320 where the Ca^2+^-binding proteins have either an Ala, Ser, or Thr residue (Figure S1*C*).

### *Substitution of Asp320 with Ala abolishes XoxF1 function with La*^*3+*^ in vivo

To test the necessity of the additional aspartate residue for Ln-dependent function of XoxF1 from *extorquens* AM1, we designed expression constructs to produce the wild-type protein and an Asp320 to Ala320 substitution variant, subsequently referred to as XoxF1 D320A. Substitution of Asp320 with Ala mimics the corresponding residue in MxaFI, the large subunit of MxaFI-type MDH in this microorganism. We chose to express wild-type and variant MDH-encoding genes via the constitutive *M*_*tac*_ promoter to bypass the complex regulatory network involved in *mxa* and *xox1* gene expression. We anticipated that expression from *M*_*tac*_ would be reduced compared to native *P*_*mxa*_ and *P*_*xox1*_ expression levels and the corresponding enzyme activities would be lower *in vivo*. As such, we tested for construct functionality in the Δ*xoxF1* Δ*xoxF2* double mutant strain, which retains a genomic copy of *exaF* (Fig. 3*A*). ExaF exhibits relatively low MDH activity with Ln^3+^, allowing the Δ*xoxF1* Δ*xoxF2* strain to slowly grow (∼ 15% the rate of wild-type cells) using methanol as the substrate (Fig. 3*B*, 3*C*), but only if Ln^3+^ are added to the growth medium. When XoxF1 was produced in the Δ*xoxF1* Δ*xoxF2* background and cells were grown with methanol and La^3+^, the culture growth rate increased by 25% and the culture growth yield increased by 67% compared to the empty plasmid control strain, Δ*xoxF1* Δ*xoxF2*::*M*_*tac*_*-empty* [*p*<0.001 by one-way analysis of variance (ANOVA)] (Fig. 3*B*, 3*C*). These results indicated the plasmid produced functional XoxF1. The strain producing XoxF1 D320A, on the other hand, grew at the same rate as the control strain and reached a similar final culture yield, suggesting XoxF1 D320A was not functional in this condition.

**FIGURE 3.**
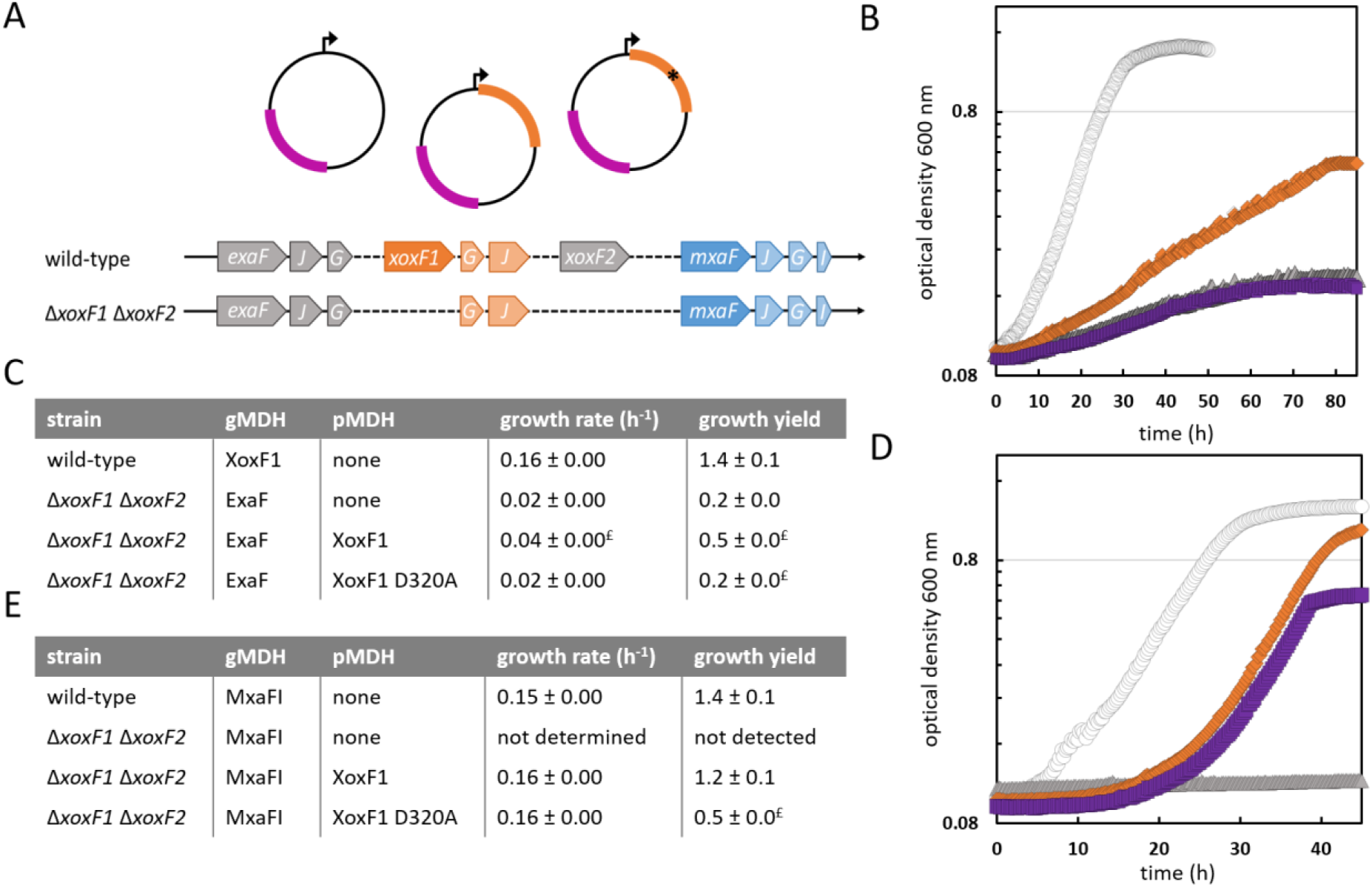
Expression of *xoxF1* and *xoxF1* D320A impacts growth on methanol in a metal-dependent manner. *A*, MDH genetic context for the wild-type and Δ*xoxF1* Δ*xoxF2* mutant strains with representation of *M*_*tac*_ expression plasmids used in these studies: empty plasmid (no insert); *xoxF1* (orange); *xoxF1* D320A (orange with black star). All constructs confer kanamycin resistance (magenta marker). *B, D*, growth curves of the Δ*xoxF1* Δ*xoxF2* MDH mutant strain carrying expression plasmids producing XoxF1 or XoxF1 D320A, respectively. Open circles, wild-type::*M*_*tac*_*-empty*; gray triangles, Δ*xoxF1* Δ*xoxF2*::*M*_*tac*_*-empty*; orange diamonds, Δ*xoxF1* Δ*xoxF2*::*M*_*tac*_*-xoxF1*; purple squares, Δ*xoxF1* Δ*xoxF2*::*M*_*tac*_*-xoxF1* D320A. Cultures were grown in minimal medium containing 20 µM CaCl_2_ with 125 mM methanol as the growth substrate. *B*, with addition of 2 µM LaCl_3_. *D*, without addition of 2 µM LaCl_3_. Growth curves are representative of a minimum of 12 biological replicates from at least two independent experiments. Replicate data points were within 5%. *C, E*, growth rates and growth yields for the wild-type and Δ*xoxF1* Δ*xoxF2* mutant strains with *M*_*tac*_ expression plasmids in the presence (*C*) and absence (*E*) of 2 µM LaCl_3_. gMDH refers to the genome-encoded ADH catalyzing methanol oxidation, if known. pMDH refers to the plasmid-encoded MDH; none, no MDH is encoded in the plasmid. Errors shown for growth rates and growth yields are RSME and standard deviation, respectively, for a minimum of 12 biological replicates from at least 2 independent experiments. ^£^ indicates a change from the Δ*xoxF1* Δ*xoxF2*::*M*_*tac*_*-empty* strain of statistical significance at *p* < 0.001 by one-way ANOVA.

### XoxF1 and XoxF1 D320A allow for equivalent growth on methanol with Ca^2+^

XoxF is required for expression of *mxa*, and, by implication, MxaFI production (51). The Δ*xoxF1* Δ*xoxF2* double mutant strain retains the *mxa* operon encoding MxaFI MDH, but it cannot grow on methanol without the addition of Ln^3+^ because it cannot produce XoxF protein (36, 51). We observed growth on methanol of the Δ*xoxF1 ΔxoxF2* strain without adding Ln^3+^ when we complemented the cells with either the *M*_*tac*_*-xoxF1* or *M*_*tac*_*-xoxF1* D320A construct (Fig. 3*D*). These results provided additional confirmation of *xoxF* expression by these constructs. Growth of double-mutant cells containing *M*_*tac*_*-xoxF1* produced enough XoxF1 to allow for a growth rate and yield similar to the wild-type strain (Fig. 3*E*). Previous work had shown that recombinant XoxF1 purified in the absence of Ln^3+^ exhibits poor activity and is insufficient to support growth with methanol as the sole MDH (52); however, that study did not establish whether XoxF1 bound Ca^2+^. Here, complementation by *M*_*tac*_*-xoxF1* D320A revealed the enzyme variant was able to execute its regulatory role. The final yield of the Δ*xoxF1* Δ*xoxF2*::*M*_*tac*_*-xoxF1* D320A culture was reduced by 58% compared to the wild-type::*M*_*tac*_*-empty* strain [*p*<0.001 by one-way ANOVA] (Fig. 3*E*). To assess whether catalytic function of the XoxF1 D320A variant was responsible for the growth defect, we conducted MDH activity assays with purified enzymes.

### Asp320 is required for catalytic function of XoxF1 MDH with Ln^3+^

Growth augmentation was observed for the Δ*xoxF1* Δ*xoxF2*::*M*_*tac*_*-xoxF1* strain when provided with La^3+^, indicating that XoxF1 was catalytically active. In contrast, analogous cells producing XoxF1 D320A showed no increase in their growth upon La^3+^ addition indicating the variant was inactive. To confirm this conclusion, 1.5-L cultures of Δ*xoxF1* Δ*xoxF2* producing either enzyme were grown in minimal methanol medium with La^3+^ and the XoxF1 and XoxF1 D320A enzymes were purified from cell-free extracts by IMAC. Sodium dodecyl sulfate-polyacrylamide gel electrophoresis (SDS-PAGE) demonstrated the successful enrichment and relative purity for both enzymes (Fig. 4*A*). XoxF1 and XoxF1 D320A were desalted and MDH activity was measured via the phenazine methosulfate (PMS)-mediated reduction of 2,6-dichlorophenol indophenol (DCPIP) (36, 53). XoxF1 was found to be active, though the specific activity with saturating substrate (*V*_max_) was only ∼50% of what we had observed in an earlier study (Fig. 4*C*) (13). This result suggested the XoxF1 used here was not fully loaded with La^3+^. An equal amount or up to 6-fold greater level of XoxF1 D320A lacked detectable activity, suggesting the enzyme did not bind La^3+^ (Fig. 4*C*). We previously reported that XoxF1 was not reconstituted by La^3+^ (13). Nonetheless, we tested whether addition of equimolar LaCl_3_ affected the assay of XoxF D320A (in case the variant enzyme could weakly bind La^3+^, or if the metal was lost during purification and/or desalting); however, no MDH activity was observed.

**FIGURE 4.**
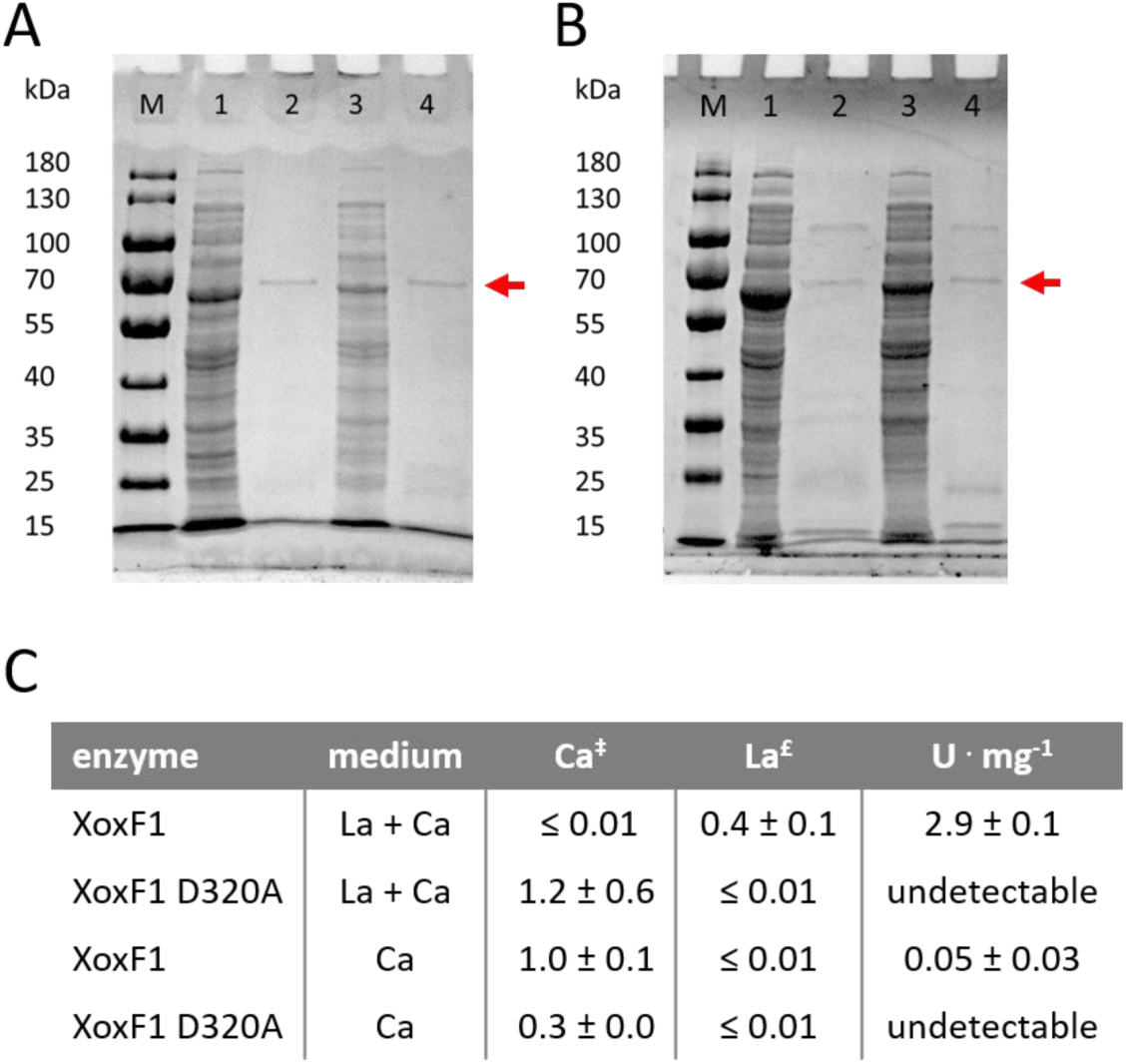
XoxF1 D320A is inactive and does not coordinate La. *A, B*; XoxF1 and XoxF1 D320A enriched from cultures of the Δ*xoxF1* Δ*xoxF2* mutant carrying an expression plasmid to produce the desired protein and grown in minimal medium with 0.5% methanol with (*A*) or without (*B*) 2 µM LaCl_3_. Enzyme fractions were analyzed by SDS-PAGE for enrichment of XoxF1 and XoxF1 D320A (predicted *M*_r_ of 63 kDa, indicated by red arrow (13)). M, protein standard marker; 1, cell-free extracts containing XoxF1; 2, 4 µg of XoxF1; 3, cell-free extracts containing XoxF1 D320A; 4, 3 µg of XoxF1 D320A. *C*, MDH specific activity measurements and metal content. MDH assays were conducted using saturating methanol substrate, with 1 unit of activity defined as 1 µmole of DCPIP reduced per min, and 4 µg of XoxF1 or 3-18 µg of XoxF1 D320A from enrichments with La^3+^; 4 µg of XoxF1 or 10-100 µg of XoxF1 D320A from enrichments without La^3+^. Values are the average of six replicates from two independent experiments with standard deviations shown. Undetectable indicates no DCPIP reduction was observed. Two independent samples of each enzyme variant were deconstructed in hot nitric acid for metal determination by ICP-AES for Ca (^‡^) and ICP-MS for La (^£^). ICP-AES was used for Ca determination because of lower background measurements compared to ICP-MS. Values are reported as moles of metal per mole subunit of enzyme. -, below background level.

Substitution of Asp320 with Ala in XoxF1 approximates the coordination environment of the MxaFI active site. We wondered, therefore, if this amino acid substitution could effectively convert XoxF1 from a Ln^3+^-dependent MDH to a Ca^2+^-dependent enzyme. Phenotypic studies of Δ*xoxF1 ΔxoxF2*::*M*_*tac*_*-xoxF1* D320A cells showed this strain was able to grow on methanol without addition of Ln^3+^, suggesting the variant was active with Ca^2+^. To investigate this possibility further, the Δ*xoxF1* Δ*xoxF2* double mutant producing either XoxF1 or XoxF1 D320A was grown in minimal methanol medium without addition of exogenous La^3+^. The IMAC-purified XoxF1 and XoxF1 D320A samples (Fig. 4*B*) were examined for their MDH activities. XoxF1 exhibited detectable activity, as also observed in a previous report (note that the variance among our measurements was relatively high, but all measured activities were low) (52). The XoxF1 D320A variant enzyme purified from the same culture condition exhibited no detectable MDH activity. The combined assay results for the XoxF1 D320A variant suggest that the Asp to Ala substitution rendered the enzyme inactive.

These results show that the single D320A amino acid substitution is not enough to convert XoxF1 into an efficient Ca^2+^-dependent MDH, and they suggest the observed growth on methanol in the absence of Ln^3+^ for Δ*xoxF1 ΔxoxF2*::*M*_*tac*_*-xoxF1* D320A cells was due to MxaFI MDH activity. In addition, the low activity observed for XoxF1 purified without La^3+^ raises the question of whether the enzyme can coordinate Ca^2+^ when the Ln-switch is not induced.

### Metal content of XoxF1 and XoxF1 D320A

Purified XoxF1 and XoxF1 D320A produced in cultures grown with and without La^3+^ were analyzed for metal content using inductively coupled plasma–(ICP)-mass spectrometry (MS) for La^3+^ or ICP-optical emission spectroscopy (OES) for Ca^2+^ quantification (Fig. 4*C*). XoxF1 purified from cells grown in medium with La^3+^ was 39% loaded with La^3+^, corresponding with the MDH specific activity observed in this study. The partial metal loading observed in this study correlates with our previous work where we observed a ∼2-fold higher *V*_max_ for XoxF1 when the enzyme was completely loaded with metal (13). In contrast to the wild-type enzyme, XoxF1 D320A purified from the same growth medium had only trace amounts of La^3+^ (Fig. 4*C*), corroborating the importance of D320 to Ln^3+^ binding by this protein. Both the wild-type and variant enzymes purified from cells grown without added La^3+^ contained trace amounts of La^3+^ likely from glass or reagent contamination, even though all glassware was acid-washed and plastic tubes and bottles were used when possible.

Although XoxF1 is Ln^3+^ dependent and expression of its gene is tightly regulated by the Ln-switch, the reported low MDH activity for XoxF1 purified from culture without added Ln^3+^ suggested it may have partial function with Ca^2+^ (52). We detected similarly low MDH activity for XoxF1 in this study and ICP-OES analysis showed the enzyme was completely loaded with Ca^2+^ (> 97%) (Fig. 4*C*). When La^3+^ was added to the culture medium, however, Ca^2+^ was not detectable in XoxF1, indicating a strong loading preference for the former metal seemingly to the exclusion of the latter for the wild-type enzyme. The XoxF1 D320A variant did not exhibit the same metal discrimination; it was loaded with Ca^2+^ regardless of whether or not La^3+^ was included in the growth medium. Although the D320A substitution does not negatively impact Ca^2+^ coordination, the enzyme is inactive. These results suggest the single amino acid substitution remodels the active site environment enough to disrupt catalysis of methanol oxidation.

### Ln-coordinating Asp is required for efficient ExaF EDH function

To date, all Ln-ADHs fall within the confines of XoxF-type MDHs and ExaF-type EDHs. In this study we have provided structural, *in vivo*, and purified enzyme biochemical studies showing the Ln-coordinating Asp is required for XoxF1 MDH function with La^3+^. To address the question of necessity of this residue in ExaF-type EDHs, we generated expression constructs to produce wild-type ExaF and its D319S variant. ExaF D319S parallels XoxF1 D320A, where substitution of Asp by Ser at position 319 approximates the active site of ExaA from *P. aeruginosa* (Fig. 5*A* and 1*B*), the Ca^2+^-dependent EDH that is most similar to wild-type ExaF with an available crystal structure (27, 54–56). Expression constructs were transformed into the ADH-4 mutant strain of *M. extorquens* AM1, which has clean deletions of the four known ADH-encoding genes that allow for growth with methanol or ethanol: *mxaF, xoxF1, xoxF2*, and *exaF*. Previously, we reported that the ADH-4 mutant strain was unable to grow in culture tubes with ethanol as the substrate. Using 48-well microplates in this study, however, we observed early poor growth with ethanol in the presence or absence of La^3+^. Because ExaF exhibits high catalytic efficiency using ethanol as the substrate (8), we anticipated that expression of its gene from the *M*_*tac*_ promoter would allow for cell growth if the enzyme produced was functional. The ADH-4 mutant strains producing ExaF or ExaF D319S were tested for growth in minimal ethanol medium with and without La^3+^ addition (Fig. 5*B*). The construct producing ExaF complemented the ADH-4 mutant strain when using La^3+^-containing growth medium, with a culture growth rate and yield that were ∼41% and ∼80% of the wild-type strain harboring the empty plasmid (Fig. 5*C*). The reduction in growth yield may have been due to ethanol evaporation from the growth medium as the cultures required an additional 15 h to reach maximal culture density in this condition. Without the addition of La^3+^, ADH-4::*M*_*tac*_*-exaF* grew marginally better than the ADH-4::*M*_*tac*_*-empty* control strain. The growth rate increased ∼3-fold, but the culture did not attain even a 2-fold increase from the initial low density. Combined, these results showed that ExaF did not oxidize ethanol efficiently in this condition, as expected (Fig. 5*D*). In comparison, ADH-4::*M*_*tac*_*-exaF* D319S grew similarly with and without addition of La^3+^, exhibiting marginally increased growth relative to the ADH-4::*M*_*tac*_*-empty* control strain (Fig. 5*B* and 5*D*). As observed for the ADH-4::*M*_*tac*_*-exaF* without addition of La^3+^, cultures did not achieve even a 2-fold increase from the starting density, indicating the enzyme variant was inefficient for ethanol oxidation regardless of metal availability. Importantly, the growth rate and yield of ADH-4::*M*_*tac*_*-exaF* with addition of La^3+^ were significantly greater than that of the ADH-4::*M*_*tac*_*-empty* and ADH-4::*M*_*tac*_*-exaF* D319S strains showing successful complementation by the construct with wild-type ExaF (One-way ANOVA, *p* < 0.001). These results indicate Asp319 is required for efficient ExaF function with Ln^3+^ and likely is important for Ln^3+^ coordination.

**FIGURE 5.**
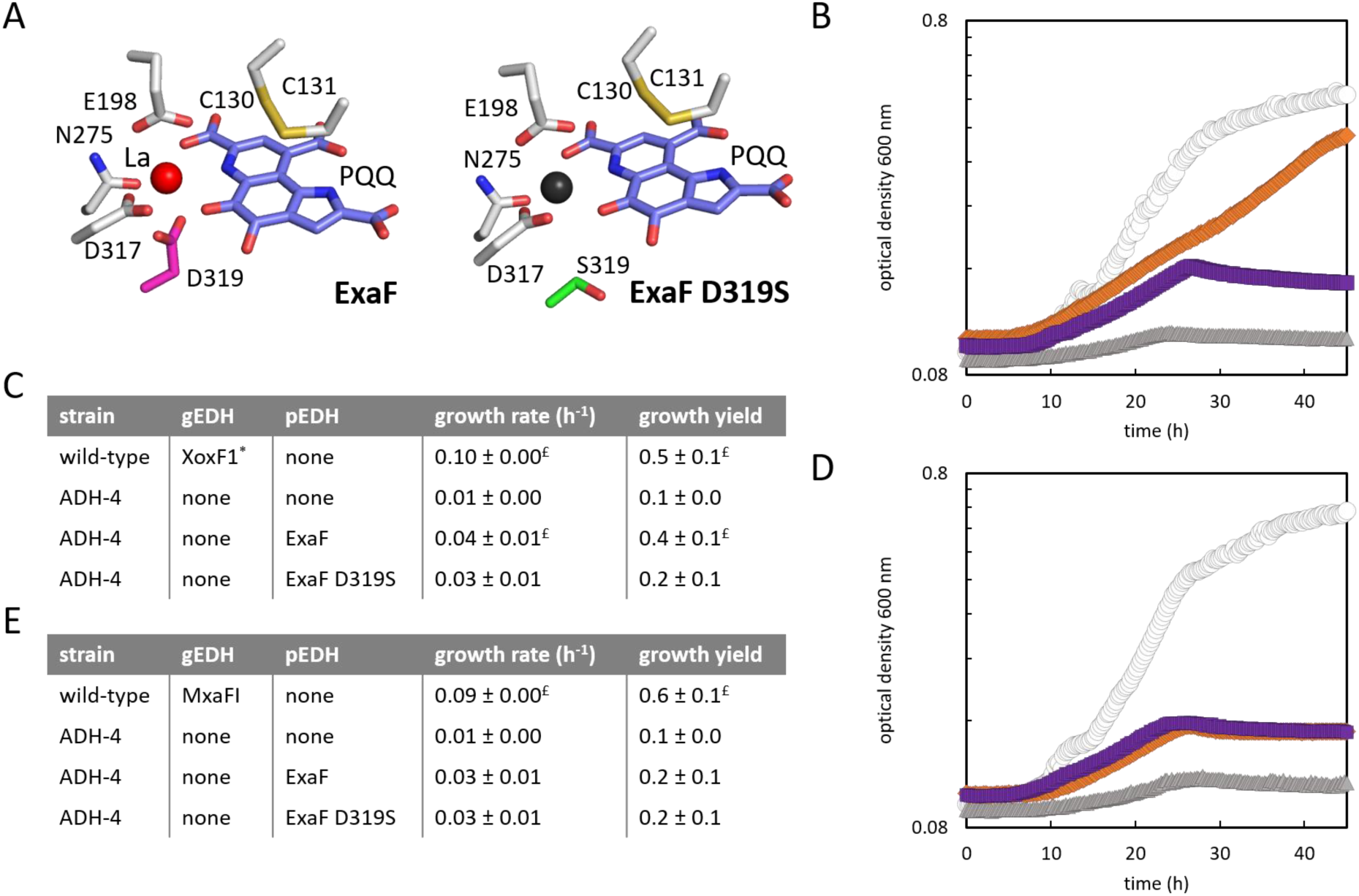
Substitution of Asp319 in ExaF EDH with Ser results in loss of function. *A*, Comparison of active site models of ExaF and ExaF D319S from *M. extorquens* AM1. PQQ, slate; La^3+^, red sphere; metal content not determined, black sphere. Ln-coordinating Asp, hot pink (ExaF); Ser substitution, green (ExaF D319S). *B*, complementation growth studies of the ADH-4 mutant with *M*_*tac*_*-exaF* and *M*_*tac*_*-exaF* D319S expression constructs. The wild-type strain carrying the empty plasmid is included as a control. Cultures were grown in minimal medium that contains 20 µM CaCl_2_ and 0.2% ethanol as the growth substrate, with (*B*) and without (*D*) addition of 2 µM LaCl_3_. Open circles, wild-type cells; orange diamonds, ADH-4 mutant cells producing ExaF; gray triangles, ADH-4 producing ExaF D319S; purple squares, ADH-4::*M*_*tac*_*-empty*. Growth curves are representative of a minimum of 18 biological replicates from four independent experiments. Replicate data points were within 5%. *C* and *E*, growth rates and growth yields for all strains included in *B* and *D*. gEDH refers to the genome-encoded enzyme catalyzing ethanol oxidation if known; none, the primary oxidation enzyme is unknown. ^*^ Proposed active EDH. pEDH refers to the plasmid-encoded EDH; none, the plasmid does not encode an EDH enzyme. Errors shown for growth rates and growth yields are RSME and standard deviation, respectively, for a minimum of 18 biological replicates from 4 independent experiments. ^£^ indicates a change from the ADH-4::*M*_*tac*_*-empty* and ADH-4::*M*_*tac*_*-exaF* D319S strains of statistical significance at *p* < 0.001 by one-way ANOVA.

## DISCUSSION

XoxF-type MDHs are members of type I eight-bladed β propeller quinoproteins (PQQ-containing). MxaFI-type MDHs and ExaF-type EDHs/ADHs fall within the same type I classification (20, 57, 58). Phylogenetic analyses have identified at least 5 major clades for XoxF-type MDHs and 9 additional clades encompassing ExaF-type EDHs/ADHs (10, 20). Yet, the number of Ln ADHs available for study are relatively few and structural data are limited. Here we report two structures of XoxF1 from *M. extorquens* AM1 (a type V XoxF MDH), one showing coordination of the La^3+^-PQQ cofactor complex and the other with only the La^3+^ atom bound. Both structures confirm coordination of La^3+^ by Asp320, as has been observed for the corresponding residue in the three previously reported XoxF MDH structures (7, 9, 41), denoting the importance of this residue for Ln^3+^ coordination and function. Comparative analysis of fully metallated subunits of XoxF1-La^3+^-PQQ and XoxF1-La^3+^-with the chain B of XoxF1-La^3+^-(61% occupied by La^3+^) indicated that Asp320 is immobile compared to the relatively flexible Trp258, Trp280, Asp318, and Arg345 sidechains. Asp320 may therefore be a key residue for recruiting Ln^3+^ to the active site of XoxF1. In addition, XoxF1-La^3+^ is the first quinoprotein structure reported without PQQ and shows that the organic cofactor is not essential for metal binding even though it provides three coordinating atoms. Incomplete occupancy of La^3+^ in chain B of this structure shows that while PQQ likely plays a stabilizing role in Ln^3+^ coordination, it appears to be a minor one. These results imply the Ln^3+^ may be loaded independently of PQQ when the Ln^3+^-PQQ complex is assembled. Additional genetic, biochemical, and structural studies are needed to gain a fuller understanding of the metal-PQQ cofactor assembly, including determination of whether the process is similar for Ca-ADH. Furthermore, dissociation of PQQ from the active enzyme to yield XoxF1-La^3+^ did not disrupt the dimeric structure of the enzyme, as speculated by Featherston *et al.* (59), showing that PQQ is not essential for maintaining dimeric interface contacts. It remains to be seen if PQQ is essential for maintaining dimeric and/or tetrameric contacts in MxaFI MDH, ExaA EDH, and other PQQ ADHs.

In this study we show that substitution of the proposed “Ln-coordinating Asp” by Ala renders the XoxF1 D320A variant unable to coordinate La^3+^, resulting in the loss of its MDH function. The parallel substitution yielding ExaF D319S results in loss of its *in vivo* function as well. Together, these results provide empirical evidence showing the necessity of the additional Asp residue in both XoxF1-type MDHs and ExaF-type EDHs and they substantiate the Ln-coordinating Asp hypothesis for determining the metal coordination and enzyme function. The identification of putative Ln ADHs by sequence alone has relied on the validity of the Ln-coordinating Asp hypothesis, which we have now corroborated with biochemical and phenotypic evidence. As a result, enzymes that have been marked as putative Ln ADHs can be investigated for Ln-utilization with a high degree of confidence, and newly discovered sequences and novel enzymes can be interrogated for the hallmark residue.

Intriguingly, the metal contents of XoxF1 and its D320A variant show that Asp320 is needed for La^3+^, but not Ca^2+^, binding *in vivo*. Insertion of Ca^2+^ into XoxF1 had been an open question since low MDH activity was reported for enzyme purified from culture medium without added Ln^3+^ (52). In this study, we corroborate those results and provide evidence that wild-type XoxF1 coordinates an equimolar ratio of Ca^2+^ when Ln^3+^ are not available. The *mxa* operon encoding MxaFI also contains genes that code for accessory proteins involved in enzyme maturation and metal insertion (39). Genes encoding a cognate cytochrome *c*_*L*_ (*xoxG*) and an essential protein of unknown function (*xoxJ*) are located in a cluster with *xoxF1*, but genes encoding a Ln^3+^ insertion system have yet to be identified (60). A separate gene cluster for lanthanide utilization and transport (*lut*), however, has been identified and characterized (61). The *lut* cluster contains several genes encoding Ln^3+^ binding proteins that also may facilitate metal insertion into XoxF1. Wild-type XoxF1 only possesses La^3+^ when purified from culture medium containing both La^3+^ and Ca^2+^, indicating a selective preference for Ln^3+^. However, we observed high levels of Ca^2+^ in the D320A variant purified from the same culture conditions. These results suggest that Asp320 may be necessary not only for Ln^3+^ coordination at the active site, but also for Ln^3+^ selectivity. One possibility to explain this observation is that the supposed “metal-free” XoxF1 binds free Ca^2+^, which is available from the growth medium for transport to the periplasm. In any case, we propose that metal selection involves the active site residues with the participation of Asp320. We also observed high Ca^2+^ content in wild-type enzyme and the D320A variant when purified from culture medium without added La^3+^. Under this condition, the Ln switch cannot occur and the *mxa* operon is expressed, including the genes encoding Ca^2+^ insertion proteins (13). It is possible that the Ca^2+^ insertion machinery encoded by the *mxa* cluster also recognizes XoxF1, however additional components are not necessary for Ca^2+^ coordination by XoxF1. More detailed knowledge of the insertion machineries is needed to fully understand how Ln^3+^ are preferentially loaded into XoxF MDH and what distinguishes Ln^3+^ insertion from Ca^2+^ insertion. XoxF1 exhibits a clear preference for Ln^3+^ as corroborated by the inactivity of XoxF1 loaded with Ca^2+^ compared to that coordinating Ln^3+^. The Ca^2+^–bound XoxF1 exhibits low MDH activity using the dye-linked assay, even though high levels of the metal are present. Thus, XoxF1 from *M. extorquens* AM1 serves as a useful representative enzyme for comparing the impacts of Ln^3+^ versus Ca^2+^ on enzyme function since it can coordinate both metals. Kinetic, mutational, crystallographic, and DFT studies with the newly available XoxF1 structure (PDB: 6OC6) will provide additional insight into how these metals affect XoxF MDH activity.

In addition to its Ln^3+^-dependent catalytic function, XoxF1 plays a role in regulation of the *mxa* and *xox1* operons. A copy of *xoxF* is required for *mxa* expression, leading to a proposed model where metal-free XoxF1 senses Ln^3+^ (36, 51). A *xoxF* suppressor mutant from the closely related *M. extorquens* PA1 is responsive to Ln^3+^, however, calling into question the essentiality of XoxF1 for Ln^3+^ sensing (62). It is worth noting that the suppressor mutations are located in the *mxbD* sensor kinase gene, whose product sits downstream of XoxF1 in the regulatory model. The resulting change to the HAMP domain of MxbD could affect signal transduction and obviate the need for XoxF1. While there is debate regarding specific details of the complex regulatory cascade, it is the structure of XoxF—not the *in vivo* MDH catalytic activity— that is crucial for regulation. In this study, we provide further evidence supporting this model using catalytically non-functional XoxF1 D320A, which allows for growth with methanol in the absence of Ln^3+^. When either XoxF1 or the D320A variant is produced without Ln^3+^ the cultures grow similarly to the wild-type strain, suggesting that *mxa* expression is similar to that in the wild-type cells. Using this condition, MxaFI catalyzes methanol oxidation, since XoxF1 does not bind Ln^3+^ and is inactive, as confirmed by our MDH assay results with pure enzymes. Our metal content analyses indicate that under these conditions XoxF1 coordinates Ca^2+^. In addition, we observed Ln^3+^-dependent growth phenotypes when producing XoxF1 D320A in the Δ*xoxF1* Δ*xoxF2* double mutant, indicating that Ln^3+^ were “sensed” by this strain. This strain does not produce a functional XoxF1 capable of coordinating Ln^3+^, reaffirming the role of the XoxF protein in regulation rather than its catalytic activity. These results suggest that XoxF1 with Ca^2+^ may be an important signal for inducing MxaFI production and further explain the binary metal loading preferences we observed by ICP-MS/OES analyses with purified enzymes.

In conclusion, our results have increased our understanding of Ln^3+^ ADH structure and function and provide two new crystal structures of XoxF1 MDH to the scientific community. These structures will aid in future endeavors to investigate Ln^3+^ and PQQ biochemistry.

## EXPERIMENTAL PROCEDURES

### Generation of MDH expression constructs

XoxF1 was produced for crystallization screens using pNG284 (containing the *P*_*xox1*_ promoter, *xoxF1* [META1_1740], and sequences encoding recombinant Tobacco Etch Virus (rTEV) protease cleavage site (63, 64) and a hexahistidine tag) in the wild-type strain of *M. extorquens* AM1 (13). To generate additional expression plasmids for enzyme production and complementation studies, PCR primers were designed with 20-40 bp overlaps between the plasmid backbone and gene inserts. For *xoxF1* expression, pNG308 was constructed by replacing the *P*_*xox1*_ promoter in pNG284 with the *M*_*tac*_ promoter and RBS_*fae*_ (65). We used pHC61 as the DNA template for the promoter with the RBS_*fae*_ sequence included in the forward primer. *M*_*tac*_ is constitutive in *M. extorquens* AM1. For *exaF* expression, pNG305 was generated using pNG308 as the DNA template for the backbone and pNG265 as the template for the *exaF* insert. The empty plasmid control, pNG311, was generated by linearizing pNG305 via PCR using a forward and reverse primer targeting the rTEV cleavage site and RBS_*fae*_ respectively. Each primer was designed with an additional 20 bp of homology to its primer partner allowing for recircularization of the now empty plasmid. All plasmids were assembled by Gap Repair assembly as described (13, 66). Amino acid substitutions were made using the Q5 Site-directed Mutagenesis kit (New England Biolabs, Ipswich, MA, USA) to generate pNG309 and pNG307 for expression of *xoxF1* D320A and e*xaF* D319S, respectively. All plasmids were verified by Sanger sequencing (Genewiz, South Plainfield, NJ, USA) and transformed into *M. extorquens* AM1 by tri-parental mating (13) or electroporation (67). Primers used for construct generation and mutagenesis are listed in Table S2.

### Enzyme expression and purification

All glassware used for protein production cultures was pre-cleaned of Ln by using it to grow the Δ*mxaF* strain on MP minimal medium (68) with 0.5% methanol. Cultures were grown with shaking at 200 rpm at 30 °C on an Innova 2300 platform shaker (Eppendorf, Hamburg, Germany) to maximal culture density. Flasks were cleaned and autoclaved, and this process was repeated until the Δ*mxaF* strain no longer grew above the initial optical density at 600 nm (OD_600_), as described (13). For enzyme or variant protein enrichment, we scaled up to a 1.5 L culture volume using 2.8 L shake flasks and grew until reaching densities of OD_600_ 1.5-6. Single colonies of strains were inoculated into 2 mL of minimal medium containing 2% succinate and 50 µg/mL kanamycin in 14 mL polypropylene culture tubes (Fisher Scientific, Waltham, MA, USA), then grown to mid-exponential growth phase with shaking at 200 rpm and 30 °C on an Innova 2300 platform shaker. Large-scale cultures producing XoxF1 and XoxF1 D320A were grown with 0.5% methanol and 2 µM LaCl_3_ or 20 µM LaCl_3_ for XoxF1 crystallization. Cells were harvested by centrifugation using a Sorvall RC6+ centrifuge (Thermo Fisher Scientific, Waltham, MA) at 21,000 x *g* at 4 °C for 10 min. Extracts were prepared as described using an OS Cell Disrupter set at 25,000 psi (Constant Systems Limited, Low March, Daventry, Northants, United Kingdom) (8). IMAC was used to purify enzymes as described (8). Enzyme enrichments were validated by SDS-PAGE analyses and desalted by buffer exchange into 25 mM Tris-HCl, 150 mM NaCl, pH 8.0, before measuring MDH activity.

### Protein crystallization

The Ln-PQQ bound protein crystals were obtained by mixing 0.65 μl of ∼2.5 mg/ml XoxF1 (reconstituted with equimolar La^3+^) and 0.65 μl of reservoir solution. The sitting drop reservoir contained 50 μl of 0.2 M ammonium chloride and 20% polyethylene glycol (PEG) 3350. Thin needles were briefly cryo protected in 25% glycerol and 75% reservoir solution prior to freezing in liquid nitrogen. For the Ln-only bound protein crystals, we mixed 0.65 μl of ∼2.5 mg/ml XoxF1 (reconstituted with equimolar La^3+^) and 0.65 μl of reservoir solution. The sitting drop reservoir contained 50 μl of 10% propanol, 0.1 M HEPES, pH 7.5, and 20% PEG 4000. A large plate-shaped crystal was frozen directly in liquid nitrogen.

### Diffraction data collection, structure determination, and analysis

X-ray diffraction data were collected at the Advanced Photon Source LS-CAT beamline 21-ID-F. Datasets were processed with xds (69), with merging and scaling done using aimless (70). Phases were solved with Phenix Phaser (71) using MDH from *M. fumariolicum* SolV (4MAE) as the starting model. Model building and refinement were conducted in COOT (72) and Phenix (73). Statistics for the datasets are listed in Table S1. Structure figures were created with UCSF Chimera (74).

### Metal quantification

Enzyme samples were deconstructed in 14 mL polypropylene tubes by heating at 90°C for 1 h in 20% nitric acid. These samples were clarified of debris by centrifugation at 21,000 x *g* for 20 min at room temperature using a Sorvall Legend X1R centrifuge (Thermo Fisher Scientific, Waltham, MA). One mL of supernatant was diluted with MilliQ water to a volume of 12 mL in a new polypropylene tube. For La^3+^ quantification, samples were sent to the Laboratory for Environmental Analysis (Center of Applied Isotope Studies, University of Georgia) for analysis by ICP-MS. Ca^2+^ quantification of enzymes was determined using a Varian 710-ES ICP-AES (Agilent, Santa Clara, CA, USA). ICP-AES resulted in lower background levels compared to ICP-MS for Ca^2+^. A MilliQ water blank and desalting buffer were analyzed as controls for background La^3+^ and Ca^2+^ contamination.

### Methanol dehydrogenase activity assays

MDH activity was measured by following the PMS-mediated reduction of DCPIP (ε_600_ = 21 mM^−1^ cm^−1^) (8, 13, 75) as described (53). The following notations are included for the assay preparation and execution: DCPIP and PMS were prepared in amber 1.5 mL Eppendorf tubes and kept on ice. Enzyme (3-100 µg) was incubated with 10 µL of 250 mM methanol or water (for no substrate controls) for 2 min at 30 °C before initiating the assay by addition of 180 µL of the dye mixture, prepared immediately beforehand at room temperature (8, 13). Little to no endogenous methanol-independent reduction of DCPIP was observed when following these modifications. Heat-inactivated enzyme controls used protein that was denatured at 95 °C for 10 min before the assay.

### Complementation in liquid culture

Single colonies of strains were inoculated into 2 mL Ln-free MP minimal medium (68) with 2% succinate, and grown in 14 mL polypropylene culture tubes (Fisher Scientific, Waltham, MA, USA) to mid-exponential growth phase with shaking at 200 rpm on an Innova 2300 platform shaker, at 30 °C. Cells were harvested by centrifugation at 1,000 x *g* for 10 min at room temperature using a Sorvall Legend X1R centrifuge. Spent culture medium was removed and cell pellets were gently resuspended in 1 mL of Ln-free MP to wash the cells. This process was repeated a second time, after which the cells were resuspended to an OD_600_ of 6 to generate starting inocula for growth studies. Growth phenotypes were compared using a BioTek EpochII microplate reader (BioTek, Winooski, VT) (13). Briefly, 10 μL of inoculum was added to 640 μL of growth medium with: 0.5% methanol or 0.2% ethanol, 50 µg/mL kanamycin, with or without 2 μM LaCl_3_. Cultures were shaken at 548 rpm at 30 °C and the OD_600_ was monitored at 15 min intervals for 48-96 h. OD_600_ measurements were fitted to an exponential model for microbial growth using CurveFitter (http://www.evolvedmicrobe.com/CurveFitter/). Growth curves were reproducible for a minimum of 12-18 distinct biological replicates from 3-4 independent experiments. Growth rates were calculated using a minimum of 40 data points. Lines of best fit were determined by an exponential model with a semi-log plot of OD_600_ vs. time. R^2^ values for all lines of best fit were > 0.99 for methanol-grown cultures and 0.98 for ethanol-grown cultures.

## ACKNOWLEDGMENTS

We thank both C. Suriano and A. Locke for their assistance in generating expression constructs and performing growth curves. This material is based upon work supported by the National Science Foundation under Grant No. 1750003 to N.C.M.-G. and N.M.G., and CHE-1516126 to R.P.H. and J.H. M.F. was supported by the University of Otago Health Sciences Postdoctoral Fellowship [HSCDPD1703]

## TABLES

**TABLE 1.** Bacterial strains and plasmids used in this study.

| strain or plasmid | description | reference |
| --- | --- | --- |
| **strains** |  |  |
| *Escherichia coli* |  |  |
| DH5α | electrocompetent cloning strain | Invitrogen |
| S17-1 | conjugating donor strain | (76) |
| *Methylobacterium extorquens* |  |  |
| AM1 | wild-type; rifamycin-resistant derivative | (77) |
| $\Delta mxaF$ | deletion mutant | (78) |
| $\Delta xoxF1 \Delta xoxF2$ | double deletion mutant | (51) |
| ADH-4 | $\Delta mxaF \Delta xoxF1 \Delta xoxF2 \Delta exaF$ quadruple deletion mutant | (13) |
| **plasmids** |  |  |
| pRK2013 | helper plasmid, IncP *tra* functions, Km<sup>r</sup> | (79) |
| pHC61 | Km<sup>r</sup>, *M<sub>tac</sub>-empty* | (65) |
| pNG284 | Km<sup>r</sup>, *P<sub>xoxI</sub>-xoxF1*, TEV cleavage site, hexahistidine tag | (13) |
| pNG286 | Km<sup>r</sup>, *P<sub>xoxI</sub>-exaF*, TEV cleavage site, hexahistidine tag | this study |
| pNG265 | Km<sup>r</sup>, *P<sub>xoxI</sub>-exaF*, Xa cleavage site, hexahistidine tag | (8) |
| pNG311 | Km<sup>r</sup>, *M<sub>tac</sub>-empty*, TEV cleavage site, hexahistidine tag | this study |
| pNG308 | Km<sup>r</sup>, *M<sub>tac</sub>-xoxF1*, TEV cleavage site, hexahistidine tag | this study |
| pNG309 | Km<sup>r</sup>, *M<sub>tac</sub>-xoxF1* D320A, TEV cleavage site, hexahistidine tag | this study |
| pNG305 | Km<sup>r</sup>, *M<sub>tac</sub>-exaF*, TEV cleavage site, hexahistidine tag | this study |
| pNG307 | Km<sup>r</sup>, *M<sub>tac</sub>-exaF* D319S, TEV cleavage site, hexahistidine tag | this study |

**TABLE S1:** X-ray data collection, reduction and refinement statistics.

| **XoxF1** | **La bound** | **La-PQQ bound** |
| --- | --- | --- |
| **Data collection** |  |  |
| Beamline | LS-CAT<br>21-ID-F | LS-CAT<br>21-ID-F |
| Wavelength (Å) | 0.979 | 0.979 |
| Space Group | P2<sub>1</sub>2<sub>1</sub>2<sub>1</sub> | P2<sub>1</sub>2<sub>1</sub>2 |
| Unit cell a, b, c (Å) | 58, 88, 202 | 101, 102, 103 |
| $\alpha, \beta, \gamma$ (°) | 90, 90, 90 | 90, 90, 90 |
| <sup>a</sup> Resolution (Å) | 48.61 – 2.80 Å<br>(2.95-2.80) | 34.06 – 3.10 Å<br>(3.31-3.10) |
| <sup>a</sup> Unique reflections | 25,837 (3,340) | 18,610 (3,366) |
| <sup>a</sup> Redundancy | 3.9 (3.4) | 4.3 (4.3) |
| <sup>a</sup> Completeness (%) | 97.7 % (88.2) | 94.3 (95.3) |
| <sup>a,b</sup> R<sub>merge</sub> | 0.088 (0.456) | 0.238 (0.522) |
| <sup>a</sup> I/σI | 9.3 (2.0) | 5.0 (2.3) |
| <sup>a,c</sup> CC<sub>1/2</sub> | 0.994 (0.789) | 0.932 (0.730) |
| **Refinement** |  |  |
| <sup>d</sup> R<sub>work</sub> / R<sub>free</sub> | 0.246 / 0.300 | 0.194 / 0.248 |
| Ramachandran favorable (%) | 92.4 | 93.0 |
| Ramachandran outliers (%) | 0.2 | 0 |
| Rotamer outliers (%) | 0 | 0 |
| Clashscore | 11.5 | 6.0 |
| R.m.s. deviation in bond lengths (Å) | 0.004 | 0.003 |
| R.m.s. deviation in bond angles (°) | 0.687 | 0.599 |
| Average B factor (Å<sup>2</sup>) | 62.3 | 19.5 |
| PDB ID | 6OC5 | 6OC6 |

<sup>a</sup> Highest resolution shell is shown in parentheses.
<sup>b</sup> $R_{\text{merge}} = \sum_{hkl} \sum_i |I_i - \langle I \rangle| / \sum_{hkl} \sum_i I_i$ , where $I_i$ is the intensity of the $i^{\text{th}}$ observation, $\langle I \rangle$ is the mean intensity of the reflection and the summations extend over all unique reflections (hkl) and all equivalents (i), respectively.
<sup>c</sup> CC<sub>1/2</sub> is the correlation coefficient of the half datasets.
<sup>d</sup> $R_{\text{work}} = \sum_{hkl} |F_{\text{obs}} - F_{\text{calc}}| / \sum_{hkl} F_{\text{obs}}$ , where $F_{\text{obs}}$ and $F_{\text{calc}}$ is the observed and the calculated structure factor, respectively. $R_{\text{free}}$ is the cross-validation R factor for the test set of reflections (5% of the total) omitted in model refinement.

**TABLE S2.**
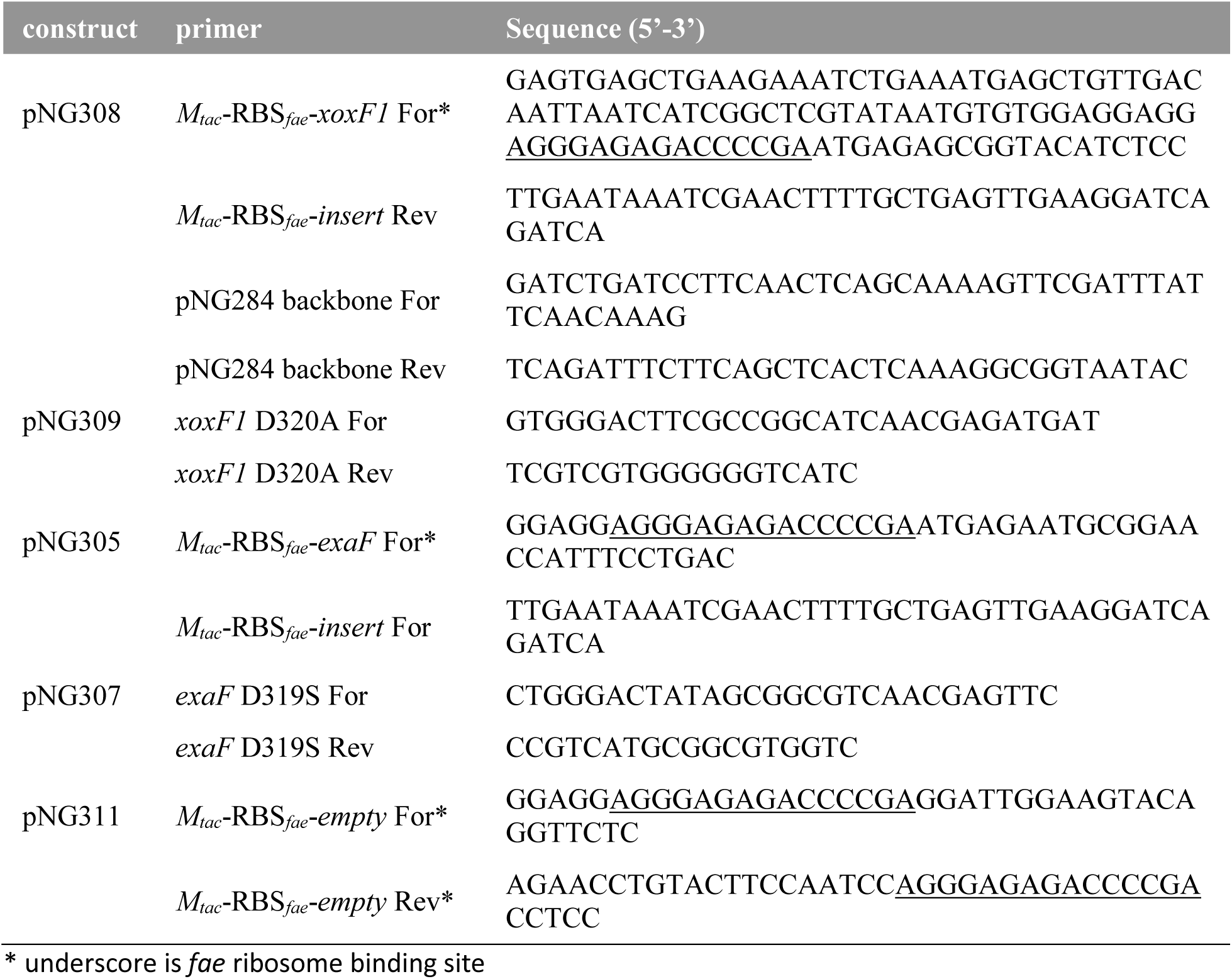
DNA oligos used for PCR amplification and plasmid assembly.

## FIGURES

**FIGURE S1.**
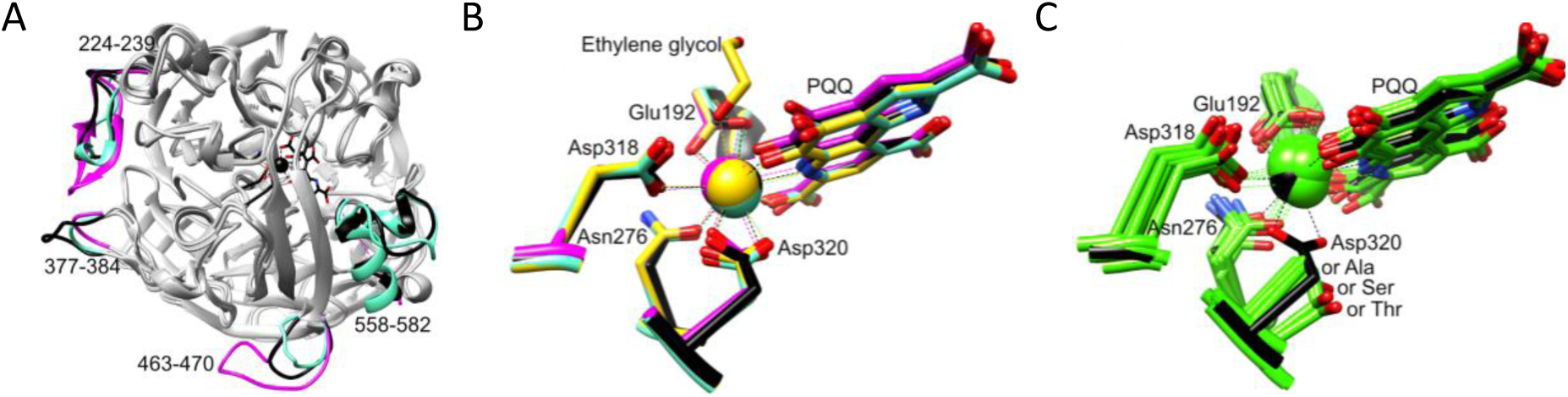
Cα alignments of MDH structures. *A*, Comparison of the overall fold for *M. extorquens* AM1 XoxF1 (black, PDB ID 6OC6, La^3+^-PQQ bound), *M. buryatense 5G* XoxF (magenta, 6DAM, La^3+^-PQQ bound), and *M. fumariolicum SolV* XoxF (cyan, 6FKW, Eu^3+^ bound). Loop numbering and active site is shown for 6OC6. *B*, Active site comparison of the same three structures, with the addition of a Ce^3+^ (and ethylene glycol) bound form of *M. fumariolicum SolV* XoxF (gold, 4MAE). *C*, Active site of *M. extorquens* AM1 XoxF1 in black compared to Ca^2+^ (or Mg^2+^)-bound MDH structures in green. The amino acid corresponding to Asp320 in XoxF1 is Ala in *Hyphomicrobium denitrificans* (2D0V (82)), *Paracoccus denitrificans* (1LRW (83)), *Methylobacterium extorquens* (1W6S (80)), *Methylococcus capsulatus Bath* (4TQO (44)), *Methylophaga aminisulfidivorans* (5XM3, the only structure with Mg^2+^ instead of Ca^2+^ (45)), and *Methylophilus methylotrophus* (1G72 (84)); Thr in *Comamonas testosterone* (1KB0 (28)) and *Pseudomonas putida HK5* (1KV9 (85)); or Ser in *Pseudomonas aeruginosa* (1FLG (81)).

